# Frequency and power of human alpha oscillations drift systematically with time-on-task

**DOI:** 10.1101/263103

**Authors:** Christopher S.Y. Benwell, Raquel E. London, Chiara F. Tagliabue, Domenica Veniero, Joachim Gross, Christian Keitel, Gregor Thut

**Affiliations:** Psychology, School of Social Sciences, University of Dundee, Dundee, UK; Centre for Cognitive Neuroimaging, Institute of Neuroscience and Psychology, University of Glasgow, Glasgow, UK; Department of Experimental Psychology, Ghent University, 9000 Ghent, Belgium; CIMEC – Center for Mind/Brain Sciences, Università degli Studi di Trento, Trento, Italy; Institut für Biomagnetismus und Biosignalanalyse, Westfälische Wilhelms-Universität, Malmedyweg 15, 48149 Münster, Germany

**Keywords:** EEG, alpha, non-stationarity, oscillations, power, frequency

## Abstract

Oscillatory neural activity is a fundamental characteristic of the mammalian brain spanning multiple levels of spatial and temporal scale. Current theories of neural oscillations and analysis techniques employed to investigate their functional significance are based on an often implicit assumption: In the absence of experimental manipulation, the spectral content of any given EEG- or MEG-recorded neural oscillator remains approximately stationary over the course of a typical experimental session (~1 hour), spontaneously fluctuating only around its dominant frequency. Here, we examined this assumption for ongoing neural oscillations in the alpha-band (8:13 Hz). We found that alpha peak frequency systematically decreased over time, while alpha-power increased. Intriguingly, these systematic changes showed partial independence of each other: Statistical source separation (independent component analysis) revealed that while some alpha components displayed concomitant power increases and peak frequency decreases, other components showed either unique power increases or frequency decreases. Interestingly, we also found these components to differ in frequency. Components that showed mixed frequency/power changes oscillated primarily in the lower alpha-band (~8-10Hz), while components with unique changes oscillated primarily in the higher alpha-band (~9-13Hz). Our findings provide novel clues on the time-varying intrinsic properties of large-scale neural networks as measured by M/EEG, with implications for the analysis and interpretation of studies that aim at identifying functionally relevant oscillatory networks or at driving them through external stimulation.

## 1. Introduction

Rhythmic neural activity is ubiquitous across multiple spatial scales of the nervous system and across species (Buzsáki et al., 2013). Since the invention of electroencephalography (EEG; see Berger, 1929), reliable associations have been revealed between distinct cognitive states/functions and frequency-specific EEG activity in humans (Kahana, 2006; Ward, 2003). This has led to proposals that oscillatory activity represents a fundamental mechanism underlying information encoding and transfer between distinct brain regions and across temporal scales (Bonnefond et al., 2017; Fries, 2005, 2015; Salinas and Sejnowski, 2001; Schyns et al., 2011; Siegel et al., 2012; Varela et al., 2001).

Oscillatory alpha-band (~8-13 Hz) activity represents one of the most prominent features of the EEG in the waking brain. A substantial body of evidence links this activity to processes such as perception (Benwell et al., 2017; Chaumon and Busch, 2014; Furushima et al., 2017; Samaha and Postle, 2015; van Dijk et al., 2008; VanRullen, 2016), attention (Foxe and Snyder, 2011; Keitel et al., 2018b; Thut et al., 2006; Worden et al., 2000) and memory (Bonnefond and Jensen, 2012; Jokisch and Jensen, 2007; Klimesch, 1999, 2012). Alpha-power is inversely related to both the blood-oxygen-level dependent (BOLD) signal (Laufs et al., 2003; Magri et al., 2012; Scheeringa et al., 2016) and cortical excitability (Haegens et al., 2011; Lange et al., 2013; Romei et al., 2007). It has been linked to the functional inhibition of cortical regions responsive to information that is irrelevant for the task at hand (Jensen and Mazaheri, 2010; Klimesch et al., 2007), potentially by gating of communication from local to other cortical regions (Fries, 2015; Zumer et al., 2014). However, neural oscillations as recorded with EEG or MEG (M/EEG) can reflect a large mixture of generating processes in terms of both local micro-circuitry and large-scale networks. For instance, recent studies have revealed multiple distinct alpha generators with different functional and/or laminar profiles (Barzegaran et al., 2017; Bollimunta et al., 2011; Haegens et al., 2015; Hughes and Crunelli, 2005; Keitel and Gross, 2016b; Scheeringa et al., 2016). Hence, while band-limited oscillatory activity measured at the macroscale is typically interpreted as a single process within a unitary framework (e.g. alpha for gating), this assumption is likely an over-simplification (Clayton et al., 2017).

Most M/EEG experiments employ designs and analyses that are based on a variety of implicit assumptions regarding the activity of interest (for an overview see Gross, 2014). For instance, in the absence of experimental manipulation (i.e. during trial baselines), neural oscillators are often assumed to spontaneously fluctuate around their mean frequency and power (Furushima et al., 2017; Romei et al., 2007; Samaha and Postle, 2015), leading in the case of alpha activity to the characteristic, band-limited alpha peak in the power spectrum. However, across the course of an experimental session, tonic power changes in certain frequencies (such as an increase in alpha-band power) (Kasten et al., 2016; Mathewson et al., 2009; Mathewson et al., 2015; Simon et al., 2011) and state- and stimulus-dependent fluctuations in oscillation peak frequency have been reported (Babu Henry Samuel et al., 2018; Haegens et al., 2014; Nelli et al., 2017), even during two minute resting-state EEG recordings (Cohen, 2014; for an overview see Mierau et al., 2017). These non-stationarities of oscillatory activity are likely of theoretical importance but are rarely accounted for when analysing experimental data.

Here, we analysed the degree of non-stationarity of two fundamental oscillatory characteristics – frequency and power – in the alpha-band over the course of a typical EEG experimental session. We found a systematic decrease in individual alpha peak frequency accompanied by a concurrent increase in alpha-power. Decomposing scalp-recorded alpha-band activity into multiple statistically separate sources via independent component analysis (ICA) revealed that whereas alpha sources tended towards power increases and frequency decreases overall, only some components showed both simultaneously. We conclude that the existence of multiple non-stationary endogenous processes inherent in macroscopic alpha-band activity can be made evident by M/EEG and should ideally be taken into consideration in future research on the functional importance of large-scale M/EEG activity.

## 2. Materials and methods

We analysed combined EEG data from two experiments that were recorded using the same EEG hardware and 60 scalp electrodes, after verifying that analysing each data set on its own revealed the same pattern of results. Both experiments lasted approximately the same amount of time (~1 hour) and had similar trial structures. Since our primary aim was to investigate the degree of non-stationarity of ongoing, baseline oscillatory activity (in the absence of stimulus processing) over the course of an experimental session, we combined pre-stimulus epochs (duration = 1 sec) across both datasets in order to increase statistical power.

### 2.1. Participants

Analyses were carried out on the data from 34 individuals (15 males, 19 females, mean age: 24, min = 17, max = 33). 20 participated in a line bisection experiment (Benwell et al., 2018) and 14 participated in a luminance discrimination experiment (Benwell et al., 2017; Tagliabue et al., 2018). All participants gave written informed consent and were financially compensated for their time. The studies were approved by the Ethics Committee of the College of Science and Engineering at the University of Glasgow. The experimental sessions were carried out within the same lab in the Institute of Neuroscience and Psychology at the University of Glasgow.

### 2.2. Instrumentation and stimuli

In both experiments, different visual discrimination tasks were performed but we combined the datasets because we were primarily interested in endogenous changes in the EEG activity during the pre-stimulus trial baselines occurring irrespective of the visual stimulus and the task performed. Full descriptions of the rationale, methodology and results for each experiment are available in the original papers (Benwell et al., 2018; Benwell et al., 2017) but a brief description of each experimental protocol is provided below.

### 2.3. Experiment 1

Participants performed a computerised line bisection task in which they were asked to estimate which of two segments of a pre-bisected horizontal line was shortest. On any given trial, the line could be 1 of 3 lengths and pre-bisected at 1 of 13 horizontal locations. The stimuli were presented using the E-Prime software package (Schneider et al., 2002) on a CRT monitor with a 1280 × 1024 pixel resolution and 85 Hz refresh rate. Participants were seated 70 cm from the monitor with their midsagittal plane aligned to the centre of the screen and their chin in a chin rest. Each trial began with presentation of a black fixation cross (subtending 0.4° × 0.4° VA) which remained on the screen for 3 s followed by presentation of the pre-bisected line (0.15 s). Following the disappearance of the line, the fixation cross remained on the screen until the participant indicated which end of the line had appeared shortest to them by pressing either the left (“v”) or right (“b”) response key with their dominant right-hand (right index and middle finger respectively). Each participant completed 702 trials overall, split into 9 blocks with short breaks being permitted between blocks. The entire experiment lasted approximately 1 hour.

### 2.4. Experiment 2

Participants performed a forced-choice luminance discrimination task in which they were asked to estimate whether a briefly presented Gaussian patch was lighter or darker than a grey background. On any given trial, the Gaussian patch could take 1 of 3 stimulus intensities corresponding to 25%, 50% and 75% successful detection performance (determined in a previous behavioural titration session). The experiment also included catch trials in which no stimulus was presented. The stimuli were presented using the Psychophysics Toolbox (Brainard, 1997; Pelli, 1997) in Matlab (Mathworks, Inc. USA) on a CRT monitor with a 1280 × 1024 pixel resolution and 100 Hz refresh rate at a viewing distance of 57 cm.

Each trial began with presentation of a black fixation cross, which remained on the screen for 1.4 s, followed by presentation of the dark or light Gaussian patch (0.03 s) in the upper right visual field. The fixation cross was then displayed for 1 sec before a response prompt asked participants to judge whether the stimulus had been lighter or darker than the background (by pressing ‘1’ or ‘2’ on the numeric pad of the keyboard with their right hand). After this decision, another response prompt asked participants to rate the clarity of their perception of a four-point scale. Responses were given by pressing four different buttons on the keyboard (‘0’, ‘1’, ‘2’ and ‘3’ on the numeric pad) and the following trial began immediately after this response. Each participant completed 800 trials overall split into 10 blocks with short breaks being permitted between blocks. The entire experiment lasted approximately 1 hour.

### 2.5. EEG recording and pre-processing

During both experiments, continuous EEG was recorded with two BrainAmp MR Plus units (Brain Products GmbH, Munich, Germany) at a sampling rate of 1000 Hz through 60 (Benwell et al., 2018) and 61 (Benwell et al., 2017) Ag/AgCl pellet pin scalp electrodes placed according to the 10-10 International System. Electrode impedance was kept below 10 KΩ. Note that the extra scalp electrode included in Benwell et al. (2017) was not analysed here in order to equate the montages between the two experiments. Subsequent pre-processing steps performed on the original data of each experiment are described below. Please note that despite their differences (in terms of filter cut-offs and down sampling), an initial re-analysis revealed the same pattern of pre-stimulus effects when each data set was analysed on its own, hence showing that the reported effects do not depend on any idiosyncratic pre-processing step in these two data sets.

### 2.6. Experiment 1 pre-processing

Pre-processing was performed using a combination of custom scripts incorporating EEGLAB (Delorme and Makeig, 2004) and FieldTrip (Oostenveld et al., 2011) functions in Matlab (Mathworks, USA). Offline, continuous data were filtered for power line noise using a notch filter centred at 50Hz. Additional low (100 Hz) and high-pass (0.1 Hz) filters were applied using a zero-phase second-order Butterworth filter. The data were then divided into epochs spanning −2.5:1.5 sec relative to stimulus onset on each trial. Subsequently, excessively noisy electrodes were removed without interpolation, the data were re-referenced to the average reference (excluding ocular channels) and trials with abnormal activity were rejected using a semi-automated artefact detection procedure. An independent component analysis (ICA) was then run using the *runica* EEGLAB function (Delorme and Makeig, 2004) and components corresponding to blinks, eye movements and muscle artefacts were removed. Missing channels were then interpolated using a spherical spline method.

### 2.7. Experiment 2 pre-processing

Pre-processing steps were performed using Brain Vision Analyzer 2.0 (BrainProducts). Offline, continuous data were filtered for power line noise using a notch filter centred at 50Hz. Additional low (85 Hz) and high-pass (0.1 Hz) filters were applied using a zero-phase second-order Butterworth filter. ICA was applied to identify and remove eye blinks and muscle artifacts. The data were segmented into epochs of 5 s starting −2.5 sec before stimulus onset and down sampled to 250 Hz. All epochs were then visually inspected and removed if contaminated by residual eye movements, blinks, strong muscle activity or excessive noise.

### 2.8. Split-half FFT analysis

In order to investigate changes in power and frequency over time, the artifact-free single-trial data were split into two separate datasets corresponding to the 1^st^ and 2^nd^ halves of the experiment for each participant and re-epoched from −1:0 sec relative to stimulus onset (i.e. 1 sec single-trial baselines). The 1 sec single-trial baselines were then multiplied by a Hamming window, zero padded and fast Fourier transformed in order to retrieve power spectra (frequency resolution = 0.1 Hz) which were averaged over trials separately for the 1^st^ and 2^nd^ half of the experiment. This yielded one power spectrum per split-half and per EEG electrode. The peak alpha frequency (frequency with highest power between 8-13 Hz) and amplitude were then extracted from the electrode with the highest mean alpha (8-13 Hz) power within each participant separately for each half of the experiment. Paired-samples t-tests were employed to test for systematic changes in both frequency and power over time. Individual changes in alpha frequency and power were calculated by subtracting the values from the 1^st^ half of the experiment from those in the 2^nd^. Hence, positive values indicated an increase in frequency/power and negative values indicated a decrease. A Spearman’s correlation analysis was performed to assess any relationship between the individual changes in alpha frequency and power over time.

### 2.9. EEG time-frequency transform

Next, we compared the frequency-power relationship at both short (within-trial) and long (across trials) time scales. Fourier-based spectro-temporal decomposition of the artifact-free single-trial data was performed using the ft_freqanalysis function (frequency-domain wavelet convolution; method: ‘mtmconvol’) from the FieldTrip toolbox (Oostenveld et al., 2011), yielding a complex-valued time-frequency plane for each trial. A temporal resolution was maintained by decomposing overlapping 0.5 sec segments of trial time series, consecutively shifted forward in time by 20 ms. Data segments were multiplied with a Hanning taper and then zero-padded to achieve a frequency resolution of 1 Hz across the range of 1:40 Hz. Power values were calculated as the squared absolute values of the complex Fourier Spectra. The data were then re-epoched from −1:1 sec relative to stimulus onset.

### 2.10. Instantaneous alpha frequency (Inst-AF) calculation

The instantaneous frequency is defined as the change in the phase of an oscillator per unit time (Boashash, 1992). The Inst-AF calculation was implemented using code developed by Cohen (2014). The single-trial EEG waveforms at all electrodes were filtered between 8 and 13 Hz using a zero-phase (two-pass), plateau-shaped, band-pass filter with 15% transition zones (Matlab *filtfilt* function). Phase angle time series for each electrode were then extracted from the resulting alpha waveforms by means of a Hilbert transform. The instantaneous frequency (in Hz) is defined by the temporal derivative of the phase angle time series (when scaled by the sampling rate and 2***π***). Noise-induced spikes in the Inst-AF time series were attenuated by application of several median filters of 10 different orders (i.e. corresponding to a minimum of 10 msec and maximum of 400 msec). The median of the resulting values was then taken. The Inst-AF data were then re-epoched from −1:1 sec relative to stimulus onset and averaged over samples in order to match the 20 ms resolution of the time-frequency power.

### 2.11. Analysis of the relationship between Inst-AF and alpha power

Cohen (2014) found a non-linear (inverted u-shape) relationship between Inst-AF and band-limited alpha power in two minute resting-state EEG recordings. To test the time-resolved relationship between Inst-AF and power here, each variable was z-transformed across all data points (20 ms resolution), trial baselines and electrodes within each participant. Note that only pre-stimulus baseline time points (−1:0 sec) were included for z-normalisation. An inverted u-shaped relationship was formally tested using the two-line solution (Simonsohn, 2017) which involves performing linear regression between the variables separately for ‘high’ (z-values above 0 collapsed across all samples (electrodes, trials, time points)) and low (z-values below 0) values of the x-axis variable (Inst-AF values). If a significant u-shaped relationship exists, then the coefficients for the ‘high’ and ‘low’ regressions should be of opposite sign and both individually significant. To test an additional influence of time-on-task on the relationship between Inst-AF and power, the data for each variable were also averaged over samples in each single-trial baseline (−1:0 sec) and these baseline-averaged values were also z-transformed across all trials and electrodes within each participant. We tested for both monotonic and inverted u-shaped relationships by means of an overall linear regression between baseline-averaged Inst-AF and power (monotonic) and the two-line solution described above (u-shaped).

### 2.12. Single-trial time-on-task analysis of both time-resolved Inst-AF and alpha-power

Within participant correlations were then calculated between trial order (1:*N trials*) and each EEG measure separately (Inst-AF and alpha-power) for all electrodes and time-points across the full epoch (−1:1 sec relative to stimulus onset, 20 ms resolution) using a two-tailed Spearman’s rank analysis. If at a given data-point (electrode/time), the value of the EEG measure systematically changes over time, then the Spearman’s rho values should show a consistent directionality across participants. Alternatively, if the EEG measure remains relatively stationary or fluctuates spontaneously across trials, then rho values across participants should be random (centred around 0). Hence, for both Inst-AF and alpha-power, we performed one-sample t-tests (test against 0) on the Spearman’s rho values across participants at all data points (i.e. all electrodes and time points). In order to control the familywise error rate (FWER) across the large number of comparisons, cluster-based permutation testing (Maris and Oostenveld, 2007) was employed. Based on the initial one-sample t-tests, all t-values above a threshold corresponding to an uncorrected p-value of 0.05 were formed into clusters by grouping together adjacent significant time-points and electrodes. This step was performed separately for samples with positive and negative t-values (two-tailed test). Note that for a significant sample to be included in a cluster it was required to have at least 1 adjacent neighbouring significant sample in both space (electrodes) and time. The spatial neighbourhood of each electrode was defined as all electrodes within approximately 5 cm, resulting in a mean of 6.3 (min = 3, max = 8) and median of 7 neighbours per electrode. The t-values within each cluster were then summed to produce a cluster statistic. Subsequently, this procedure was repeated across 2000 permutations to create a data-driven null hypothesis distribution using *ft_statistics_montecarlo* (Oostenveld et al., 2011). On each iteration, this function effectively switched the sign of the correlation for a random subset of the participants. One-sample t-tests of rho values against zero were then performed at each data point. After clustering significant t-values across neighbouring data points (as above), the most extreme cluster-level t-score was retained. The location of the original real cluster statistics within this null hypothesis distribution indicates how probable such observations would be if the null hypothesis were true. Hence, if a given negative/positive cluster had a cluster statistic lower/higher than 97.5% of the respective null distribution cluster statistics, then this was considered a significant effect (5% alpha level). Note that, for the sake of completeness, both pre- and post-stimulus time points were included in this analysis in order to investigate whether the effects of time-on-task held for both time periods. This was the only analysis in which we also included post-stimulus time points. In order to further assess the nature of the observed monotonic relationships, we also applied linear, quadratic and cubic fits to the trial number – alpha frequency and trial number – alpha power relationships.

### 2.13. Split-half stratification FFT analysis

We then investigated to what extent the decrease in alpha frequency and increase in alpha power over time are linked by employing a stratification analysis. We hypothesised that if the frequency decrease and power increase are manifestations of the same process, holding constant the value of one measure between the 1st and 2nd half of the experiment should also abolish the change in the other measure. For example, if the Inst-AF decrease over time is co-varying exactly with the increase in power, then when power is held constant the Inst-AF should also remain stationary (and vice versa). To test this, single-trial values of both alpha power (8:13 Hz) and Inst-AF were averaged over the baseline period (−1:0 sec) across all electrodes and the data were split into bins corresponding to the 1st and 2nd half of the experiment in each participant. The mean, variance and higher order statistics of both the 1st and 2nd half trial distributions of one of the measures (either alpha-power or Inst-AF) were equalized by sub-sampling the single trials using the *ft_stratify* function (‘histogram’ method with 100 bins) in FieldTrip (Oostenveld et al., 2011). The stratification was performed separately for alpha power and frequency. Because the stratification algorithm approximately equalizes the trial distributions and the results are not necessarily identical each time it is run, the peak alpha frequency and amplitude were extracted from the electrode with the highest mean alpha (8-13 Hz) amplitude (using the same FFT analysis as described above across the trials remaining after stratification) on every iteration of 100 runs of the analysis and averaged (separately for the 1st and 2nd half distributions) within each participant. On average, 211 trials (min = 141, max = 320) remained in each distribution after stratification of alpha power and 190 trials (min = 108, max = 267) remained after stratification of alpha-frequency. The peak alpha frequency and power values were entered into paired-samples t-tests to test for systematic changes in either frequency or power over time after approximate equalisation of the other variable.

### 2.14. Alpha-band source separation using independent component analysis

In order to further investigate the relative independence of the power increases and frequency decreases over time, we performed an analysis using independent component analysis (ICA) and current-dipole modelling to statistically separate sources of alpha-activity contributing to the EEG signal measured on the scalp. Scalp-level EEG alpha activity reflects a superposition of multiple alpha rhythms (Barzegaran et al., 2017; Keitel and Gross, 2016a). Thus the primary aim of the ICA analysis was to statistically separate independent components of alpha-activity and to investigate whether the scalp-level power- and frequency trends persisted for individual alpha components. Our tests were conducted against the null-hypothesis that across all components the power and frequency trends were net zero. Alternatively, and supported by trends at the scalp level (see Results), we expected that alpha components would show a preference towards power increases and frequency decreases. Statistical separation further allowed us to examine whether each individual component echoed scalp-level **mixed** power and frequency trends or whether some components showed **unique** increases in power over time (no concurrent frequency decrease) or **unique** decreases in frequency over time (no concurrent power increase). Finally, using this classification of components we investigated whether components possessing distinct non-stationary characteristics were spectrally dissociated.

First, artifact-free epochs were band-pass filtered at 5-15 Hz, then an ICA (runica EEGLAB function (Delorme and Makeig, 2004)) was performed on the filtered concatenated epochs using only the baseline period (−1:0 sec). For each participant, we obtained 20 “alpha” IC’s by subjecting the data to a principal component analysis prior to ICA (runica option ‘pca’ set to 20). Original unfiltered epochs were then projected through the spatial filters (channel weights) of each IC. As a next step, we identified ICs that showed reliable alpha peaks (within 8-13 Hz), as determined by an automated peak-finding algorithm based on smoothing of power spectral density (PSD) estimates with an 11-point, 3^rd^ order polynomial Savitzky-Golay filter (Corcoran et al., 2018; Keitel et al., 2018a; Savitzky and Golay, 1964). For the selected components, a single equivalent current dipole was fit using a three-layer boundary element method (BEM) template model based on the standard Montreal Neurological Institutes’s (MNI) brain template using the DIPFIT EEGLAB plug-in (Oostenveld and Oostendorp, 2002) with default options. Only dipoles with fits leaving less than 15% residual variance and that were located inside the model brain volume were selected for further analysis (see Gulbinaite et al., 2017 for a similar approach). This resulted in a total number of 327 alpha-component dipoles across participants (median per participant = 10, range = 2-16). Next, single-trial power and instantaneous frequency estimates were calculated for each component by means of a Hilbert transform (see **Instantaneous alpha frequency (Inst-AF) calculation** section above, instantaneous power = squared absolute of the analytical Hilbert signal) and averaged within each trial baseline. Within each participant, Spearman rank correlations were then calculated between trial order (1:N trials) and each measure separately (single-trial Inst-AF and alpha-power) for each component. For each individual component Spearman’s rho value, z-values and non-parametric p-values were calculated against a permutation distribution of rho-estimates based on 10000 random shuffles of trial order. At the group level, we tested for systematic shifts in power and frequency using one-sample t-tests (test against 0) on the z-values across all components. Additionally, we tested whether combined power and frequency changes clustered around a preferred tendency in a two-dimensional distribution. First, collapsing the 2D- into a circular space, we computed the phase angle of a surrogate complex number whose real part was given by the z-value indicating the power change and its imaginary part given by the z-value indicating the frequency change for each component. The phase angle distribution was then tested for non-uniformity using a Rayleigh test. Second, because this analysis neglects the magnitudes of component trends, we supplemented the Rayleigh statistic with a circular T^2^ test (Victor and Mast, 1991). In the component plane spanned by power and frequency trends, 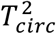 tests whether the centroid of the bivariate distribution is different from the origin (0,0). Finally, in order to investigate spectral differences, we compared peak alpha frequency values between **unique** power increase, **unique** frequency decrease and **mixed** power increase/frequency decrease components by means of independent samples t-tests.

### 2.15. Data and code availability statement

The EEG data used in this study, as well as the Matlab code to re-produce the analyses and figures, are openly available on The Open Science Framework (OSF) under the URL: https://osf.io/7sfgw/

## 3. Results

### 3.1. Baseline alpha-power and -frequency show non-stationarity over time

Figure 1A plots grand-averaged power spectra centred on the alpha band for the 1^st^ (blue) and 2^nd^ (red) half of the experimental session respectively. The peak frequency values are denoted by the vertical dashed lines. This plot suggests that over time peak alpha frequency *decreased*, whereas alpha power *increased*. Figures 1B and C plot for each participant the change in individual alpha frequency (IAF) and power over time (2^nd^ half IAF minus 1^st^ half IAF), highlighting the consistency of the effects across participants for both measures. The group median IAF was 10.35 Hz for the 1^st^ half of the experimental session(standarddeviation = 0.89 Hz, range: 8.4-11.7 Hz) and 10.1 Hz for the 2nd half (standard deviation = 0.9 Hz, range: 8.2-11.4 Hz). Although the frequency shift was small on average (0.25Hz), shifts within individuals could reach 2Hz (see Fig 1B). Paired-sample t-tests confirmed a group-level reduction in IAF (t(33) = −2.5755, p = .0147, Cohen’s d = .442) and an increase in alpha-power over time (t(33) = 3.35, p = .002, Cohen’s d = .823). Additionally, both effects persisted using proportional change measures (reduction in IAF: t(33) = −2.64, p = .0125), increase in power: t(33) = 2.8149, p = .0082). However, the individual changes in peak alpha frequency did not correlate with the individual changes in alpha power across participants (Spearman’s Rho = −0.038, p = .8305). Figure 1D plots grand-averaged topographies of alpha-power (8-13 Hz) for the 1^st^ (top) and 2^nd^ (bottom) half of the experimental session respectively, revealing similar spatial distributions but an increase in amplitude over time.

**Figure 1:**
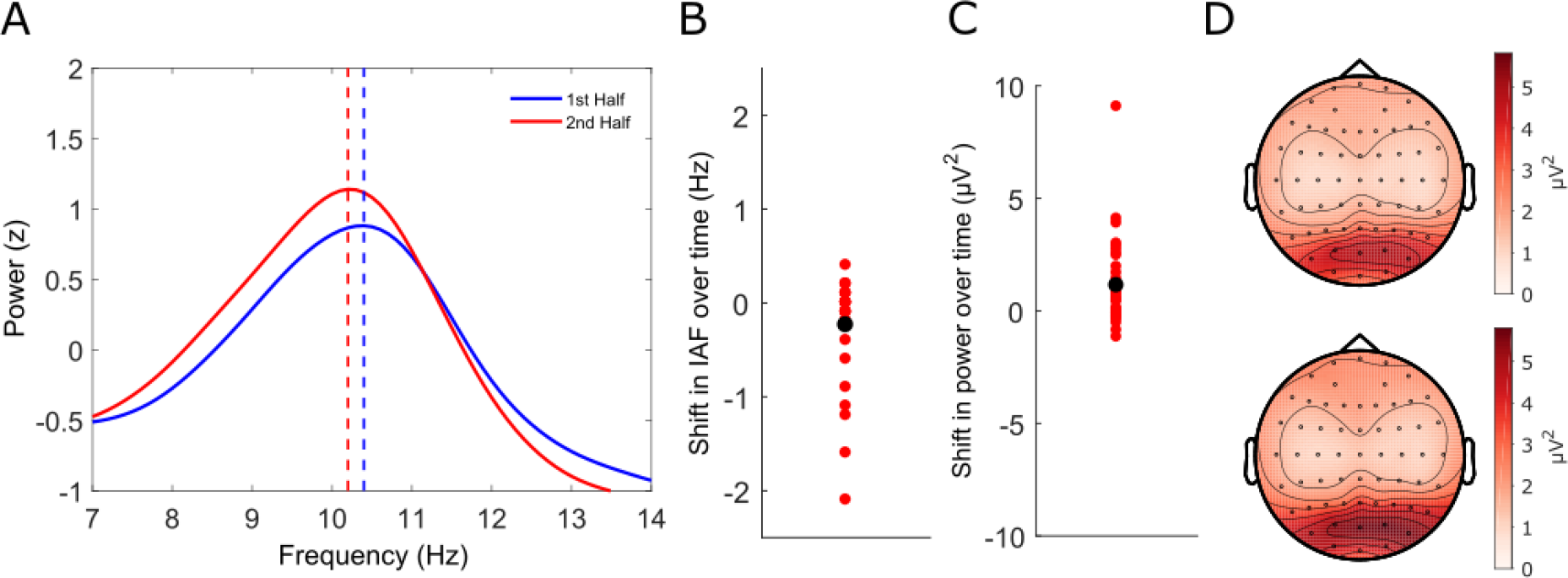
Split-half Fast Fourier Transform (FFT) analysis. **(A)** Grand-averaged frequency spectra (z-scored) in the alpha band from trial baselines for both the 1^st^ (solid blueline) and 2^nd^ (solid red line) halves of the experimental session. The peak frequency values are denoted by vertical dashed lines (blue = 1^st^ half, red = 2^nd^ half). Alpha power increased whereas peak alpha frequency decreased from the 1^st^ to the 2^nd^ half of the experimental session. **(B)** Shifts in individual alpha frequency from the 1^st^ to the 2^nd^ half of the experimental session. Red dots denote individual participants and the black dot represents the group average shift. **(C)** Shifts in individual alpha power from the 1^st^ to the 2^nd^ half of the experimental session. Red dots denote individual participants and the black dot represents the group average power change. **(D)** Grand-averaged topographies of alpha power (8-13 Hz) for the 1^st^ (top) and 2^nd^ (bottom) half of the experimental session respectively. The topographical distributions were almost identical but the increase in amplitude over time is apparent.

### 3.2. Comparing frequency-power relationship between shorter and longer time scales

Cohen (2014) found a non-linear (inverted u-shape) relationship between estimates of instantaneous alpha frequency (Inst-AF) and band-limited alpha power in two minute resting-state EEG recordings (see also Nelli et al. (2017)). Here, we replicated this finding with time-resolved data from the single-trial baseline alpha activity for which we determined alpha power and Inst-AF. Both measures were averaged within consecutive 20 ms segments for each 1-sec baseline epoch per electrode and participant separately to preserve variability on the short time scale (i.e. within trials) and then z-scored within participants. Alpha-power was then plotted against Inst-AF (see Fig 2A for a representative sample of participants, and Fig 2B for the group average). A consistent relationship between alpha power and alpha frequency across time-points of the 1-sec epoch (i.e. on a short time scale) is revealed, with increased variability and higher maximum power values close to the central alpha frequency. The inverted u-shaped relationship was formally tested using the two-line solution proposed by Simonsohn (2017). This method involved performing linear regression between Inst-AF and power separately for ‘high’ (in our case, z-values above 0) and ‘low’ (z-values below zero) values of the independent variable (Inst-AF values) for each participant. If a significant inverted u-shaped relationship exists, then the coefficients for the ‘high’ and ‘low’ regressions should be opposite in sign and both individually statistically significant. The analysis revealed a significant positive relationship between low (below 0) z-values of Inst-AF and power (t-test of coefficients across group against 0: t(33) = 8.7979, p < .0001) and a significant negative relationship between high (above 0) z-values of Inst-AF and power (t(33) = −18.8715, p < .0001), thereby formally confirming the inverted u-shaped relationship between time-resolved Inst-AF and power.

**Figure 2:**
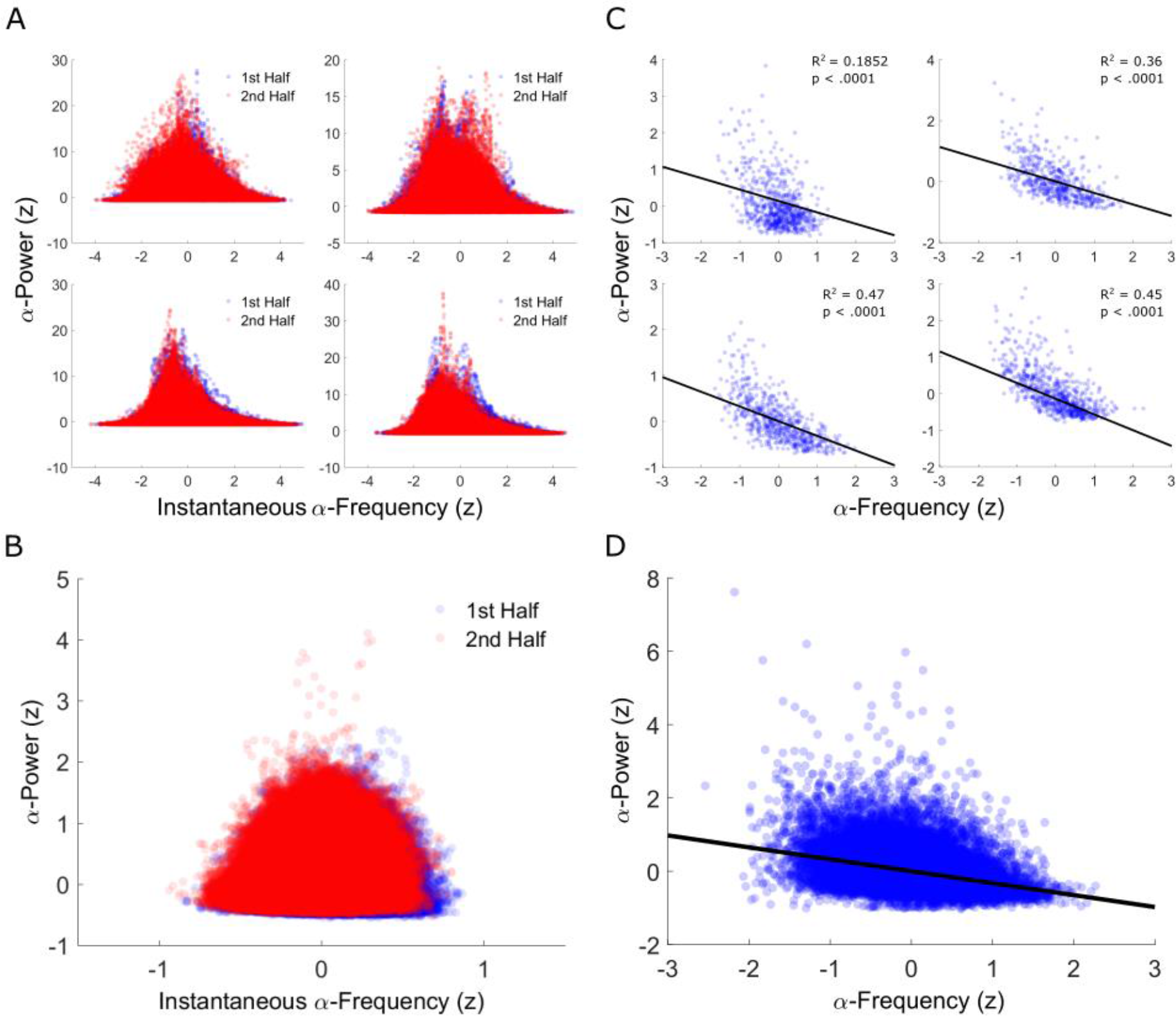
Comparing the alpha frequency-power relationship between shorter versus longer time scales (A&B versus C&D). **(A)** Scatterplots from four representative participants of alpha power values (y-axis: z-scored) as a function of instantaneous alpha frequency (x-axis: z-scored) from the 1^st^ (blue dots) and 2^nd^ halves (red dots) of the experimental session. Note that each dot represents a single electrode-time point for one participant. **(B)** Group-averaged scatterplot of short time-scalere solved alpha power values (y-axis: mean z-scores) as a function of instantaneous alpha frequency (x-axis: mean z-scores) from the 1^st^ (blue dots) and 2^nd^ halves (red dots) of the experimental session. Dots represent the group-average per single electrode-time point. Both **A** and **B** show systematic frequency-power variability with increased variability and higher amplitude of power values close to the central alpha frequency (replicating Cohen (2014)). **(C)** Scatterplots from the same 4 participants as in **A** of alpha power values (y-axis: z-scored) as a function of alpha frequency (x-axis: z-scored) when longer time-scale resolved (averaged over time-points within each single trial baseline). Hence, each dot represents a baseline-averaged value from a single-trial for one electrode, and the graph illustrates between-trial variability in pre-stimulus alpha power/frequency over time. **(D)** Group-averaged scatterplot of longer time-scale resolved alpha power values (y-axis: z-scored) as a function of alpha frequency (x-axis: z-scored). Dots represent the group-averaged baseline values for each trial and electrode. Both **C** and **D** show that emphasising variability over the longer-term time-scale(across trials) highlights an inverse relationship between alpha power and frequency, in addition to the parabolic relationship observed at the short-term scale (within trials) in **A** and **B** .

However, our split-half data binning analyses above suggest that both measures also change systematically over a longer time-scale (i.e. across trials over the course of an experiment), with alpha frequency decreasing and power increasing. Hence, we re-represented the data as frequency-power distributions after averaging Inst-AF and power values over each 1-sec baseline epoch per electrode to emphasize the variability across a longer time scale (i.e. across trials) and performed regression analysis between these values. Figure 2C-D shows the corresponding scatter plots of alpha power values against Inst-AF values for the same representative participants as in Figure 2A (see Fig 2C), along with the group average (see Fig 2D). We tested for both monotonic and nonlinear relationships between baseline-averaged Inst-AF and power and found a significant negative overall linear relationship (t(33) = −4.1123, p < .0001) but no inverted u-shaped relationship as the Inst-AF-power regression for both the low and high Inst-AF bins showed significant negative relationships (low Inst-AF values versus power: t(33) = −2.1746, p = .0369, high Inst-AF values versus power: t(33) = −6.6183, p < .0001). This confirms that introducing a longer-term time scale (across trials) results in an inverse relationship between Inst-AF and power, as opposed to the parabolic relationship observed at the short-term scale (within trials).

### 3.3. Frequency sliding and power increase confirmed with cluster-based analysis of instantaneous frequency and power

Next, we performed a time resolved analysis across 2-sec single-trial, stimulus-locked epochs spanning both baseline and post-stimulus periods (−1 to +1 sec from stimulus-onset). Post-stimulus data were included to test whether the relationship holds for both pre- and post-stimulus periods. Figure 3 displays the results of group-analyses in which we investigated whether correlation coefficients between spectral content and trial number differed systematically from zero (i.e. negative or positive) across participants. In line with the split-half analyses, the results revealed that Inst-AF was lower in the 2^nd^ as compared to the 1st half of the experimental session (Figure 3A, t-values systematically negative over entire epoch), whereas alpha-power was higher in the 2nd as compared to the 1^st^ half of the session (Figure 3D, t-values systematically positive). Their respective pre-stimulus topographic representations (averaged over −1 to −0.5 sec to avoid the possible inclusion of post-stimulus activity) are shown in Figure 3B and 3E. Figure 3C shows the relationship between trial number (x-axis) and z-scored single-trial Inst-AF (averaged over −1:−0.5 sec across all electrodes in each single-trial baseline period; y-axis) collapsed across all participants. On average, the alpha peak frequency decelerated by 0.2 Hz per hour. A Spearman’s correlation applied to this collapsed data confirmed a highly significant negative relationship (rho = −0.1131, p<.0001). The black line represents the best-fitting linear regression slope. Both linear and quadratic fits to the data were significant(95% confidence bounds for coefficients did not overlap with 0), though adding the additional term (quadratic) only minimally improved the goodness-of-fit (linear fit adjusted R-squared = 0.0130, quadratic fit = 0.0135). Adding an additional term (cubic fit) did not improve the fit (95% confidence bounds for additional coefficient overlapped with 0; adjusted R-squared = 0.0135). Figure 3F shows the relationship between trial number and z-scored single-trial alpha-power. A highly significant positive relationship was confirmed (Spearman’s Rho = 0.1222, p<.0001). Again, both linear and quadratic fits to the data were significant (95% confidence bounds for coefficients did not overlap with 0), though adding the additional term (quadratic) only minimally improved the goodness-of-fit (linear fit adjusted R-squared = 0.0105, quadratic fit = 0.0132). Adding an additional term (cubic) did not improve the fit (95% confidence bounds for additional coefficient overlapped 0; adjusted R-squared =0.0132).

**Figure 3:**
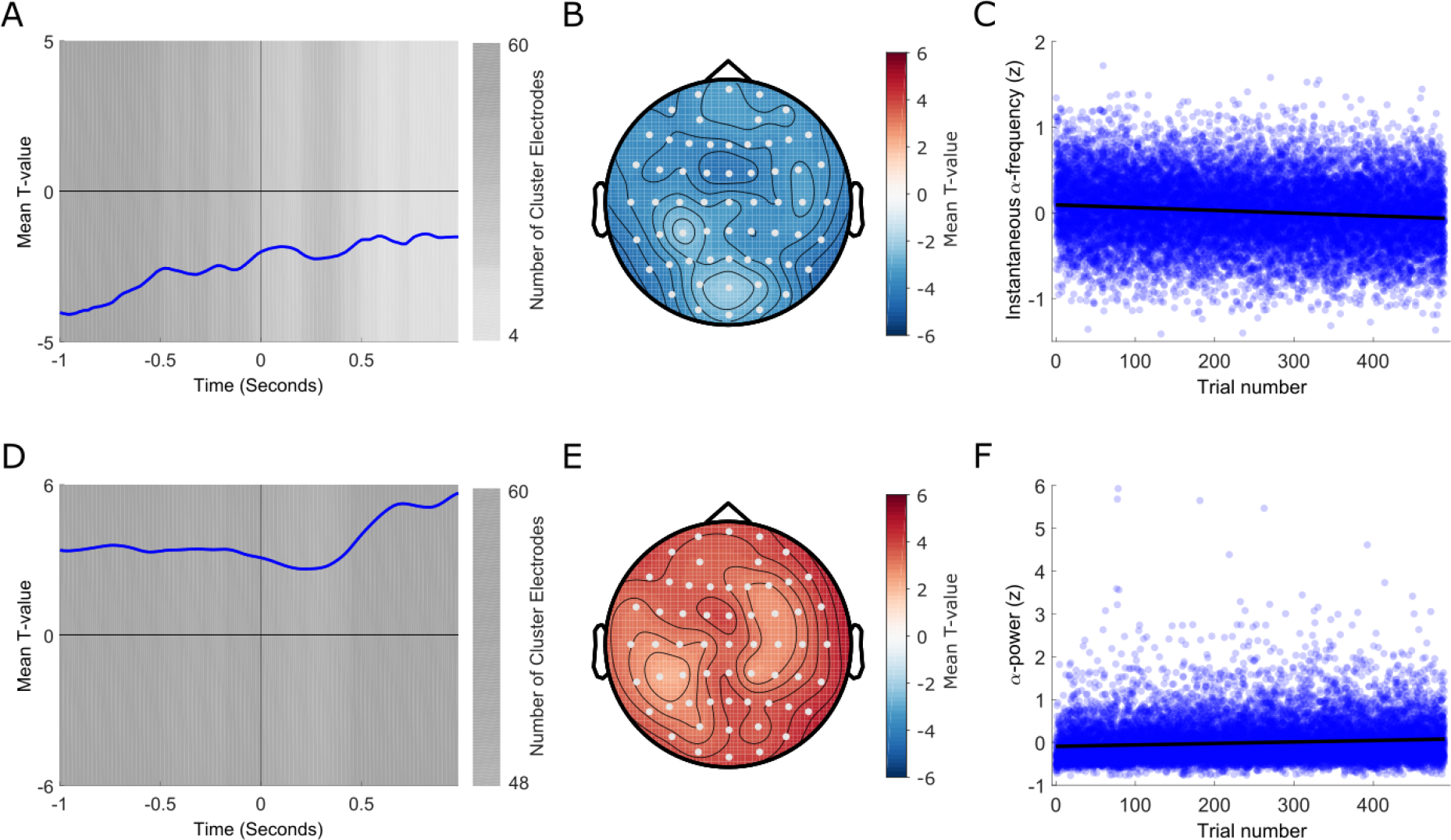
Time-resolved analysis of frequency sliding and power changes. **(A)** Trial order-instantaneous alpha frequency effect: T-values averaged over all electrodes. A negative t-value indicates a decrease in frequency over trials. Significant time-points are indexed by the grey background fill. One significant negative cluster was found which spanned the entire epoch. Hence, this analysis confirmed that alpha -frequency systematically decreased over the course of the experimental session at both pre- and post-stimulus time points. The luminance of the background fill indicates how many electrodes were included in the significant cluster at each time-point (the darker the fill, the more electrodes are included in the cluster at this time-point). **(B)** Topographic representation of the t-values averaged over the pre-stimulus (−1:−0.5 sec) portion of the significant cluster. Electrodes that were included in the cluster at least once at any time-point during this time period are highlighted in white. **(C)** Scatterplot of the relationship between trial number (x-axis) and instantaneous alpha frequency (y-axis: z-scored and averaged over −1:−0.5 sec across all electrodes in each single-trial baseline period). Note that the data for all participants are included and the number of trials included was again limited to the first 490 to equalise across participants. The solid black line represents the best-fitting regression slope. **(D)** Trial order to alpha power effect: T-values values averaged over all electrodes. A positive t-value indicates an increase in power over trials. Significant time-points are indexed by the grey background fill. One significant positive cluster was found which spanned the entire epoch. Hence, this analysis confirmed that alpha power systematically increased over the course of the experimental session at both pre- and post-stimulus time points. The luminance of the background fill indicates how many electrodes were included in the significant cluster at each time-point (the darker the fill, the more electrodes are included in the cluster at this time-point). **(E)** Topographic representation of the t-values averaged over the pre-stimulus (−1:−0.5 sec) portion of the significant cluster. Electrodes that were included in the cluster at least once at any time-point during this time period are highlighted in white. **(F)** Scatterplot of the relationship between trial number (x-axis) and alpha power (y-axis: z-scored and averaged over −1:−0.5 sec across all electrodes in each single-trial baseline period). Note that the data for all participants are included and the number of trials included was again limited to the first 490 to equalise across participants. The solid black line represents the best-fitting regression slope.

### 3.4. Frequency sliding and power increases in the alpha-band over time are driven by partially independent processes

Figures 4A-B plot the results of a power stratification analysis (i.e. when power was held constant between the 1st and 2nd split-half bins of the experiment). Despite power being held constant, the grand average peak alpha frequency still decreased from the 1st to the 2nd half of the experiment (Figure 4A) as denoted by the vertical dashed lines (blue = 1st half, red = 2nd half). Figure 4B plots the change in peak alpha frequency over time (2nd half IAF minus 1st half IAF) for each participant. A paired t-test confirmed a group-level reduction in peak alpha frequency (t(33) = −2.8497, p = .0075). Figures 4C-D plot the results of the Inst-AF stratification analysis (i.e. when Inst-AF was held constant between the 1st and 2nd split-half bins of the experiment). Again, despite Inst-AF being held constant, alpha power increased from the 1st to the 2nd half of the experiment (Figure 4C). Figure 4D plots the change in alpha power over time for each participant, which also remained significant (t(33) = 2.1759, p =.0368). Hence, when power was held constant, the decrease in peak alpha frequency over time was maintained and when Inst-AF was held constant, the increase in power over time was also maintained. Overall, this suggests that frequency sliding and power changes over time are not likely to be two manifestations of the same process but rather may index two partially independent non-stationary processes in the alpha-band.

**Figure 4:**
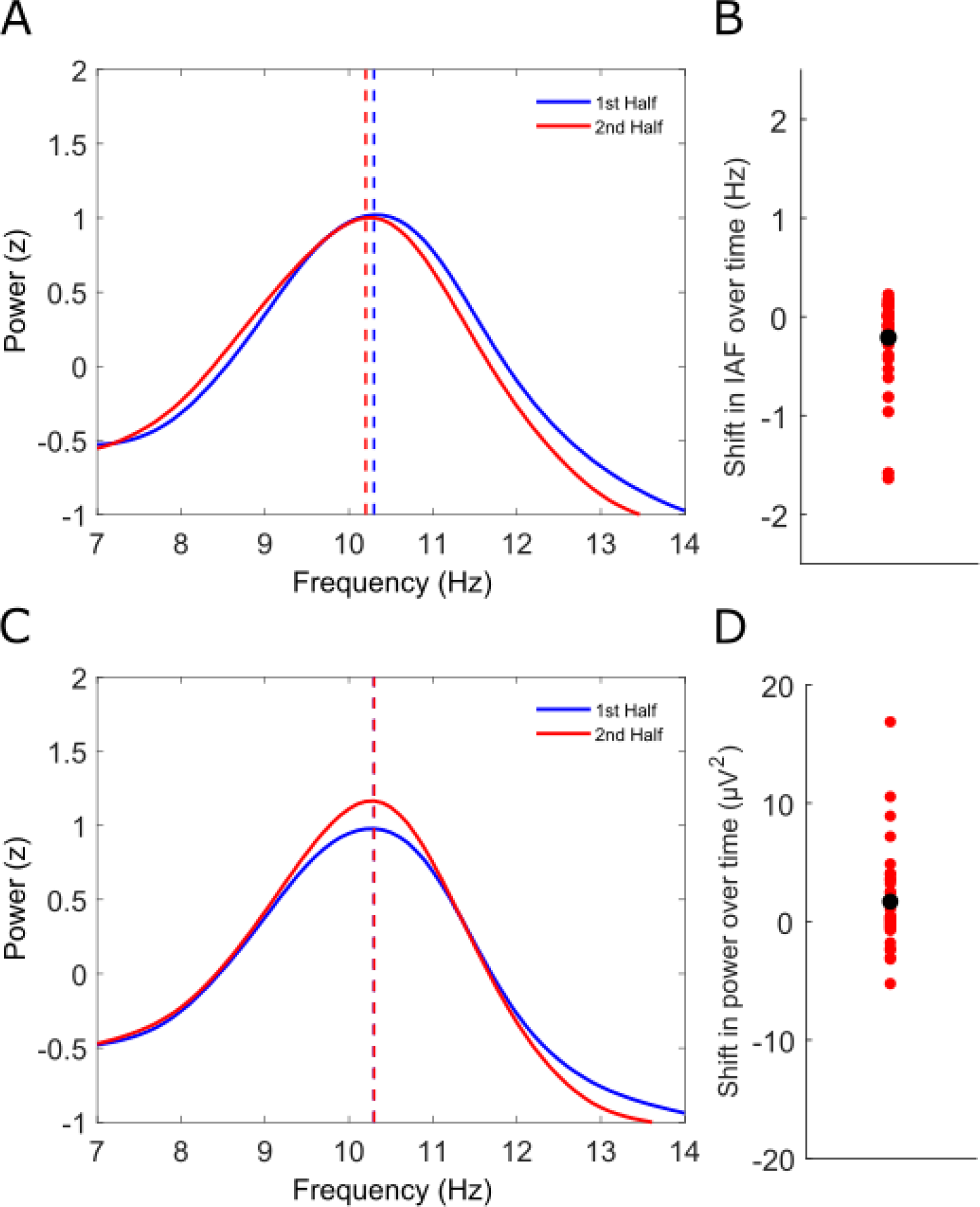
Trial stratification analysis reveals non-stationary processes are at least partially independent. **(A)** Analysis with power held approximately constant between the 1^st^ and 2^nd^ half of the experiment through trial sub-sampling: Grand-averaged frequency spectra (z-scored) in the alpha band from stratified trial baselines for both the 1^st^ (solid blueline) and 2^nd^ (solid red line) halves of the experimental session. The peak frequency values are denoted by vertical dashed lines (blue = 1^st^ half, red = 2^nd^ half). Despite alpha power remaining approximately stationary, peak alpha frequency still decreased from the 1^st^ to the 2^nd^ half of the experimental session. **(B)** Shifts in individual alpha frequency (IAF) from the 1^st^ to the 2^nd^ half of the experimental session from the power stratification analysis. Red dots denote individual participants and the black dot represents the group average shift. **(C)** Analysis with instantaneous frequency held approximately constant between the 1^st^ and 2^nd^ half of the experiment through trial sub-sampling: Grand-averaged frequency spectra (z-scored) in the alpha band from stratified trial baselines for both the 1^st^ (solid blueline) and 2^nd^ (solid red line) halves of the experimental session. The peak frequency values are denoted by vertical dashed lines (blue = 1^st^ half, red = 2^nd^ half). Despite alpha frequency remaining approximately stationary, alpha power still increased from the 1^st^ to the 2^nd^ half of the experimental session. **(D)** Shifts in individual alpha power from the 1^st^ to the 2^nd^ half of the experimental session from the instantaneous frequency stratification analysis. Red dots denote individual participants and the black dot represents the group average power change.

### 3.5. Both mixed and unique power/frequency non-stationarities contribute to sensor level effects

To further investigate the relative (in)dependence of alpha power increases and alpha frequency decreases over time, we repeated the single-trial regression analyses on ICA-derived alpha components. In line with the sensor level results, alpha components showed an overall increase in power over time (t(326) = 6.5446, p<.0001) and an overall decrease in frequency over time (t(326) = −4.9096, p<.0001). Figure 5 plots locations of equivalent dipoles and standardised regression coefficients (indexed by dot colour) for alpha independent components (masked (p < .05) at the individual component level) which showed **unique** power changes over time (5A), **unique** frequency changes over time (5B) and **mixed** power and frequency changes over time (5C). Note the dominance of increases in power over time in 5A (indexed by positive z-scores) and the dominance of decreases in frequency over time in 5B (indexed by negative z-scores). Figure 5D plots the distribution of directionalities of all component shifts over time in a 2-dimensional trend space spanning the orthogonal axes of frequency and power changes (from decrease to increase). The changes systematically cluster around a combination of power increases and frequency decreases (average angle = 335.52° +/− 13.15°, Rayleigh z = 35.2025, p<.0001), also when taking into account the magnitude of the effects (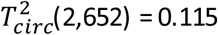, p < 1×10^−15^). The latter is illustrated in Figure 5E, which shows the 2-dimensional distribution of components with a black square indicating its centroid. A zoom-in on the central region (bottom-right panel) further depicts the centroid’s bootstrapped 95% confidence intervals (1000 samples, bias-corrected and accelerated, see Efron (1987)). Notably, both CIs exclude zero (power trend CI = [0.921 1.717]; frequency trend CI = [−0.840 −0.365]).

**Figure 5:**
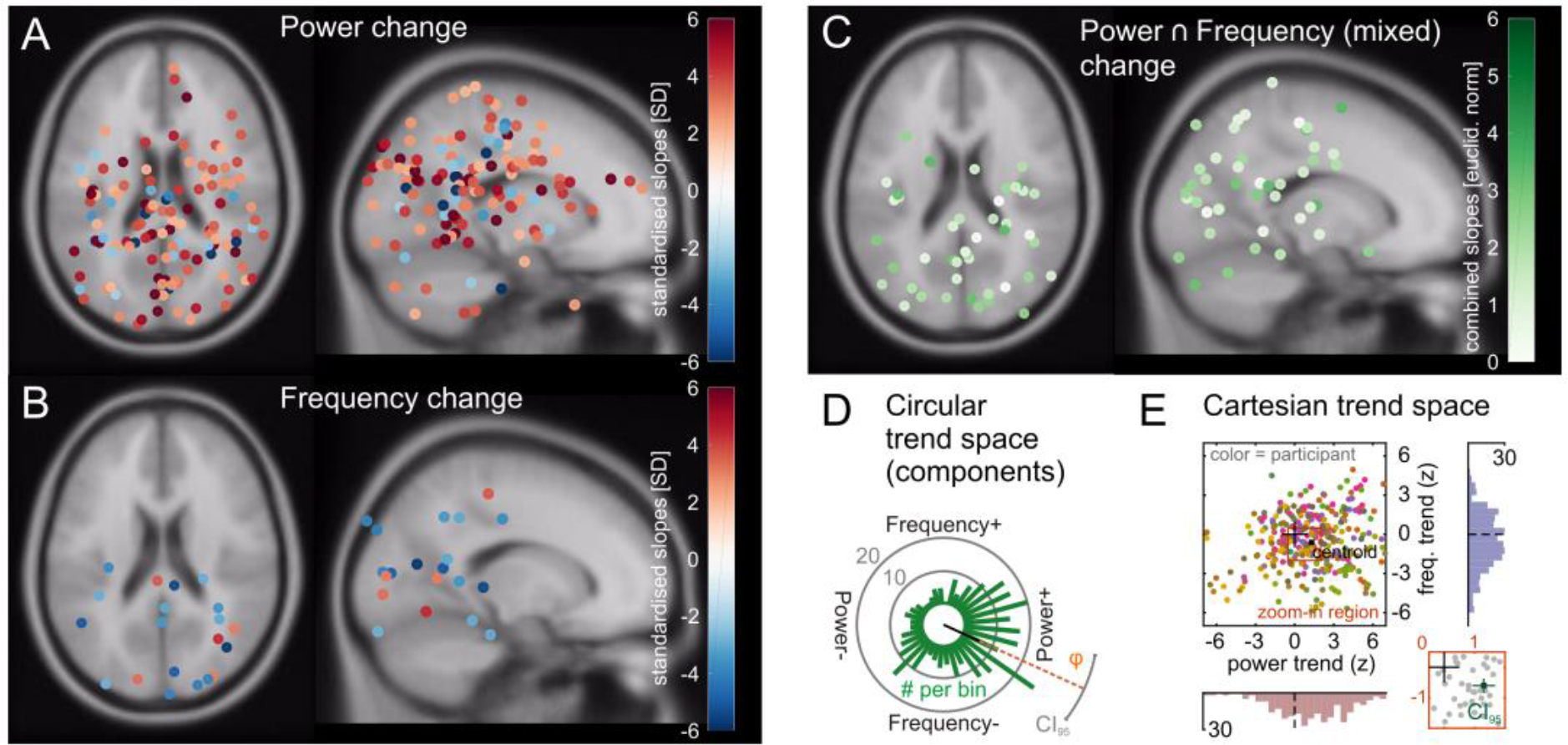
Statistical separation of alpha sources – dipole distributions and time-on-task analysis. (**A**) Dipole locations of components showing alpha power change over time, but no frequency change, in MNI source space regardless of the directionality of the change. Only component dipoles with regression coefficients corresponding to non-parametric p-values < 0.05 (uncorrected) are plotted for visualisation purposes. Note that the majority of dipoles show power increases (warm colours). (**B**) Dipole locations of components showing alpha frequency change, but no power change, in MNI sourcespaceregardless of the directionality of the change. Only component dipoles with regression coefficients corresponding to non-parametric p-values < 0.05 (uncorrected) are plotted for visualisation purposes. Note that the majority of dipoles show frequency decreases (cool colours). Also note that fewer dipoles show unique frequency change as compared with dipoles showing unique power change in A. (**C**) Source-space distribution of dipolelocations showing both alpha power and frequency changes = mixed change. The colour indicates the magnitude of the combined slopes (=Euclidean norm of power and frequency shifts). Note that more components exhibit mixed change than unique frequency changein B. Because the representation in C conceals the direction of the effects, (**D**) illustrates the distribution of directionalities in a 2-dimensional trend space spanning the orthogonal axes of frequency and power change (from decrease to increase). Bars in this circular histogram indicate the number of components (scale = grey circles) that show a similar preference towards a certain combination of change. The changes cluster around a combination of power increases and frequency decreases (black line displays mean resultant vector, grey arc shows the corresponding 95% circular confidence interval). Also note that only very few components fall into a power decrease / frequency increase region. (**E**) Distribution of alpha components by power- and frequency trend (z-scored). Data points in the main plot are color-coded to assign components to participants (same participant = same colour). The ‘+’ marks the origin (0,0). A zoom-in (bottom-right panel) on the central region of the main plot (brown square) shows bootstrapped 95%-confidence intervals for mean power- and frequency trends (green lines). Additional histograms show marginal distributions of power- (bottom) and frequency trends (right); dashed lines = zero.

Figure 6A plots grand-averaged alpha-component spectra for mixed power increase/frequency decrease components (green), unique frequency decrease components (blue) and unique power increase components (red) respectively. Mean peak frequency values are denoted by the vertical dashed lines. Figure 6B plots histograms of the corresponding distributions of individual component peak frequencies, showing that those components which both increased in power and decreased in frequency over time had significantly lower peak frequencies (at around 8-10Hz) than both unique frequency decrease components (t(192) = −6.6528, p<.0001) and unique power increase components (t(219) = −7.1865, p<.0001) (at around 10-12Hz). Unique frequency decrease components did not differ in peak frequency from unique power increase components (t(165) = 0.6686, p=.5047). These differences in spectral characteristics held even when the analysis was restricted to data from the 1^st^ quarter of trials in each participant (mixed components versus unique frequency decrease components (t(192) = −5.7509, p<.0001), mixed components versus unique power increase components (t(219) = −5.2314, p<.0001), unique frequency decrease versus unique power increase components (t(165) = 1.1790, p=.2401)). Hence, they cannot be explained by the non-stationarities themselves but rather likely represent intrinsic features of separate alpha components. Figure 6C plots power spectra in the alpha-band (z-scored) across all individual components for mixed power increase/frequency decrease components (top row), unique frequency decrease components (middle row) and unique power increase components (bottom row).

**Figure 6:**
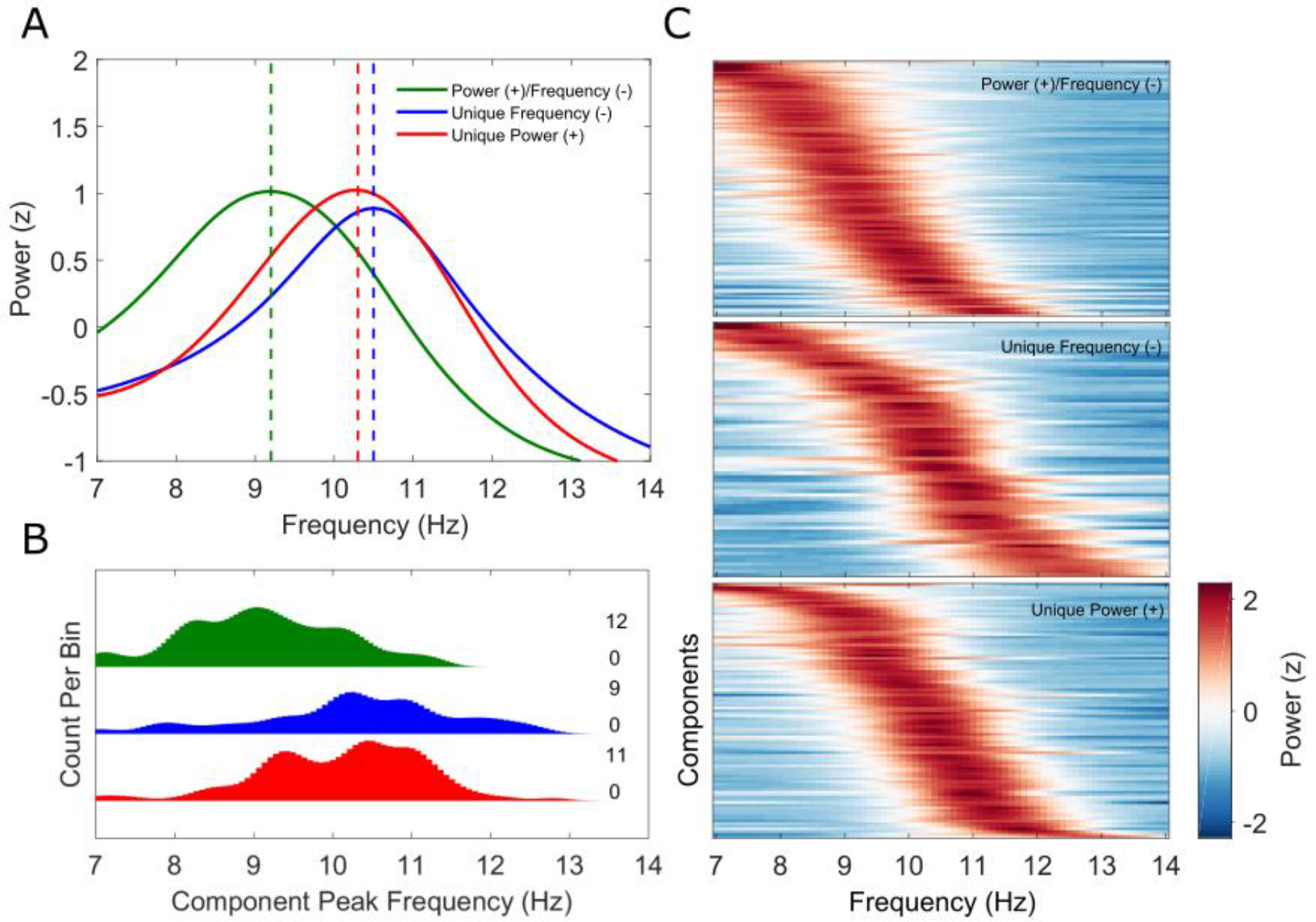
Statistical separation of alpha sources – frequency distribution analysis. (**A**) Grand-averaged power spectra (z-scored) for alpha independent components (ICs) showing both power increase and frequency decrease over time (solid green line), unique frequency decrease over time (solid blueline) and unique power increase over time (solid red line) respectively. Mean peak frequency values are denoted by the vertical dashed lines. (**B**) Distributions of individual component peak frequencies for mixed power (+)/frequency (−) (green), unique frequency (−) (blue) and unique power (+) (red) alpha ICs. Note the relatively lower peak frequency values for the mixed power (+)/frequency (−) components. Minimum and maximum frequency bin counts are plotted to the right of each distribution separately. (**C**) Power spectra of individual mixed power (+)/frequency (−) (top), unique frequency (−) (middle) and unique power (+) (bottom) alpha ICs. Each row represents an IC and colour corresponds to z-scored spectral power.

Overall, the ICA analysis suggests a mixture of underlying generating mechanisms for the sensor level effects in the alpha band, including alpha generators with distinct alpha sub-bands (see Figure 7 for a discussion of possible underlying models). The results show that a single power increase/frequency decrease oscillator model can be ruled out (Figure 7A). Instead, the effects can be partly accounted for by low alpha frequency components increasing in power which simultaneously pulls the instantaneous frequency towards lower alpha frequencies (i.e. effectively changing the ratio of low/high alpha frequency power in favour of low frequencies (see Figure 7C)). However, additional components show unique power increases and frequency decreases, in line with the model proposed in Figure 7B. Hence, multiple non-stationary processes likely contribute to the net alpha-power increase and alpha-frequency decrease over time measured at the sensor level.

**Figure 7:**
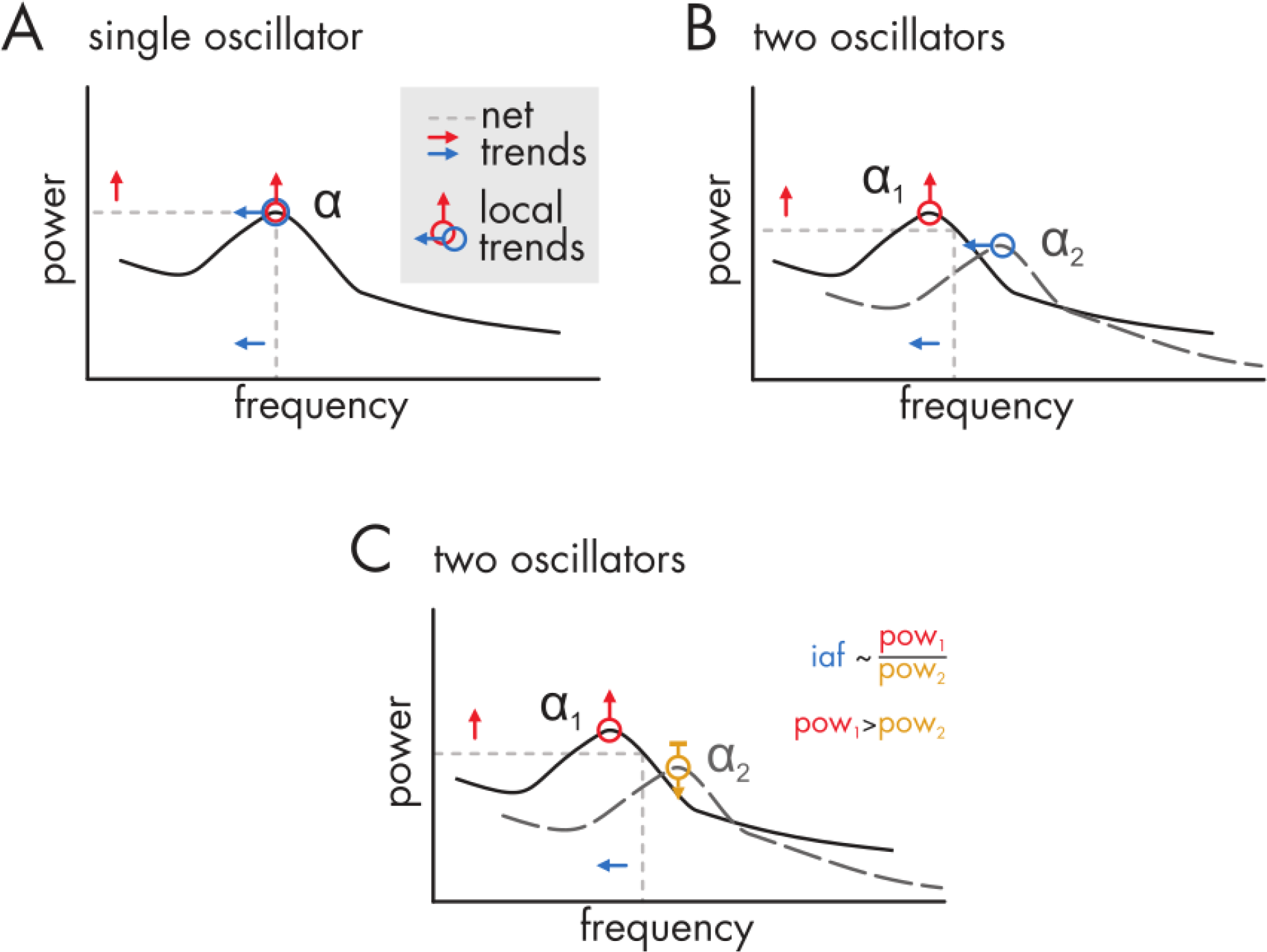
Three candidate scenarios underlying systematic long-term alpha frequency decrease and alpha power increase at the sensor level (= net trend). Black lines (straight in **A**, straight and dashed in **B** and **C**) depict schematic power spectra of underlying neural assemblies oscillating at a mean (peak) alpha frequency (with spontaneous fluctuations around the mean over trials determining peak width). Note that all scenarios produce the same net trends in changes over time (red and blue arrows on grey dashed lines). (**A**) Alpha, as measured with M/EEG, is generated by a single process that increases its power (red arrow) and reduces its frequency (blue arrow) over time. This scenario cannot account for the current results because several generative sources with distinct spectral and spatial profiles were revealed by the ICA analysis. (**B**) Our findings of independent power and frequency changes speak to the presence of multiple generative processes (black straight and dashed lines). One of them increases in power (e.g. α_1_, red arrow), while the other decreases in frequency over time (e.g. α_2_, blue arrow). (**C**) Changes in power only (red and golden arrows), of generative processes with offset peak frequencies (α_1_ and α_2_), may also contribute to the same net trends. Their power ratio can influence the estimate of individual alpha frequency becausepeaks arenot readily separable in EEG data (instead, EEG measurements capture a weighted sum of single spectra). Evidence for this scenario is provided by our finding of mixed power increase/frequency decrease components primarily in low alpha frequencies which may alter the low/high frequency alpha power ratio over time in favour of low frequency alpha which simultaneously decreases the peak frequency. Note that these scenarios are necessarily simplifications of a more complex cortical reality that may involve multiple oscillating micro-circuits (Cohen, 2017), also with non-sinusoidal properties (Cole et al., 2017; Cole and Voytek, 2017; Jones, 2016).

## 4. Discussion

The current study reveals the existence of endogenous non-stationary processes in human alpha-band EEG activity. Over the course of an experimental session (~1 hour), peak alpha-frequency decreased whereas alpha-power increased, even in the absence of external stimulation during trial baselines. Furthermore, the frequency decrease and power increase were partially independent and hence did not represent two manifestations of the same process. Below, we discuss these findings from theoretical, methodological and applied perspectives in light of current theories and approaches in electrophysiological research on oscillatory brain activity and its functions.

### 4.1. Understanding neural network activity at the macro-scale

Over recent decades, evidence has converged on a dynamic functional role of alpha oscillations in sensation and cognition beyond the seminal alpha-idling hypothesis (Pfurtscheller et al., 1996). Alpha oscillations have been implicated in the orchestration of active inhibition vs. facilitation of task-irrelevant vs. task-relevant brain regions (Foxe and Snyder, 2011; Klimesch et al., 2007; Rihs et al., 2007), of sensory sampling and its temporal resolution (Cecere et al., 2015; Samaha and Postle, 2015) and the interregional gating or communication of information (Fries, 2015; Jensen and Mazaheri, 2010; Palva and Palva, 2007; Zumer et al., 2014). Here, we reveal systematic non-stationarities in alpha power and peak frequency. The increase in alpha-power over time replicates previous studies, which have associated the effect with the level of sustained attention and fatigue (Boksem et al., 2005; Cajochen et al., 1995; Craig et al., 2012; Kasten et al., 2016; Mathewson et al., 2015; Simon et al., 2011). Accordingly, it is plausible that changes in vigilance and fatigue over the course of the experimental session may be related to the shift in alpha power and also frequency observed here. Drifts in oscillatory power within experimental sessions have also been linked to non-stationarities in psychophysical measures (Benwell et al., 2013; Benwell et al., 2018; Bompas et al., 2015; Mathewson et al., 2009) and cognitive processes necessary for the maintenance of task performance (Stoll et al., 2016). In terms of alpha-frequency, previous studies have shown long-term changes over the life-span related to aging and pathology (Aurlien et al., 2004; Klimesch, 1999; Mierau et al., 2017) as well as short-term changes in the sub-second to second range. For instance, alpha-frequency transiently increases in response to sensory stimulation and task engagement, particularly under conditions of high stimulus intensity and/or cognitive demand (Babu Henry Samuel et al., 2018; Cohen, 2014; Haegens et al., 2014; Hülsdünker et al., 2016; Maurer et al., 2014; Nir et al., 2010). Here we add the crucial observation that alpha frequency systematically *decreases* over the course of a typical experimental session in the order of minutes/hour.

While in line with previous studies reporting non-stationarities in alpha-power and frequency (on the time scale of seconds), our data provide novel insights into their potential sources, albeit at the limited spatial resolution of EEG. The macroscale activity of widespread, larger cell assemblies has been postulated to produce slower oscillations and also higher power density than local, smaller cell assemblies (Buzsaki, 2004). Therefore, an increase in power and concurrent decrease in frequency may be explained by changes in the extent of oscillating networks over time. Additionally, previous studies have linked cycle-by-cycle increases in oscillatory power with decreases in instantaneous frequency, both in rat hippocampal gamma-activity (Atallah and Scanziani, 2009) and in evoked human alpha-activity (Himmelstoss et al., 2015). The relationship has been attributed to rapid adjustments in synaptic inhibition in response to fluctuating excitatory amplitude. Although our results are partially commensurate with these interpretations, our ICA analysis suggests a mixture of generating mechanisms with distinct spectral and spatial profiles. Alpha-activity in EEG recordings has previously been dissociated in lower and upper sub-bands (Klimesch et al., 1997; Lobier et al., 2018; Shackman et al., 2010), which are thought to be differentially implicated in various cognitive and perceptual functions (ElShafei et al., 2018; Klimesch, 1999). In line with the presence of meaningful alpha sub-divisions, our results show low frequency alpha networks (~8-10Hz) to be associated with tonic power increases over time and a simultaneous decrease in peak frequency, while independent, higher alpha frequency components (~9-13Hz) showed unique decreases in instantaneous frequency or increases in power respectively. Hence, the scalp level alpha non-stationarities do not reflect a unitary rhythm but a mixture of generating mechanisms with potentially different neurophysiological and functional characteristics. These findings open up exciting possibilities to further separate out and functionally dissociate alpha networks with a view to establishing a more complete understanding of the role they play in perception and cognition (e.g. by dissociating those components which show unique frequency changes, power changes, or both). It is likely that some of the observed non-stationary alpha components are related to fluctuations in global vigilance and fatigue states (Cajochen et al., 1995; Sadaghiani et al., 2010), whilst others may be specific to engagement with cognitively or perceptually demanding tasks (Stoll et al., 2016). These distinct networks can be further disentangled by future studies to identify sources of distinct alpha non-stationarities and their functional relevance at the macro-scale.

### 4.2. Methodological and applied considerations

Systematic endogenous non-stationarity of both the frequency and power of alpha activity calls for caution in the interpretation of some frequently employed M/EEG time-frequency analysis procedures and their results. For instance, in the absence of experimental manipulation, stationarity of oscillatory characteristics is often assumed, or intrinsic (i.e. baseline) fluctuations are interpreted to represent stochastic or spontaneous processes (Caspers, 1983; Meisen, 2016; Samaha and Postle, 2015). However, band-limited EEG spectral non-stationarities occur at the sub-second timescale (Cohen, 2014; Gross, 2014; Meisen, 2016) and also in the order of minutes/hours as demonstrated in the present study.

This is of importance because apparently spontaneous fluctuations in pre-stimulus power, phase and frequency are increasingly being linked to perceptual and cognitive functions (Benwell et al., 2017; Busch and VanRullen, 2010; Furushima et al., 2017; Mathewson et al., 2009; Samaha and Postle, 2015). On average, we found changes in frequency of 0.2 Hz/hour. While this may appear to be small, and is an order of magnitude smaller than the typical alpha band width, some studies have identified brain-behaviour relationships with absolute effect sizes well within the range of the non-stationarities we observe here, hence suggesting that changes of this magnitude are of perceptual and cognitive relevance (see Samaha and Postle (2015) and Wutz et al. (2018) for reports of perceptually relevant frequency differences of ~0.02-0.04 Hz). In particular, non-stationarities should be considered for the interpretation of M/EEG-behaviour relationships when not only present in the M/EEG signal but also displayed in the employed behavioural measures, as can be the case for psychometric function threshold and slope, detection rate and reaction time (Benwell et al., 2013; Benwell et al., 2018; Doll et al., 2015; Frund et al., 2011). In this situation, a correlation between the two measures of interest may be detected simply because both measures systematically change over time (i.e. they both independently correlate with trial order across the experiment). If this is not investigated and/or accounted for, then a potentially epiphenomenal correlation between the two measures may be misinterpreted as a causal or functional relationship. Hence, it is critical that non-stationarity is considered in the interpretation of any observed brain/behaviour relationship (see Benwell et al., 2017; Mathewson et al., 2009; van Dijk et al., 2008 for examples of studies which explicitly account for temporal variations in M/EEG and/or behavioural measures). If non-stationarity is present in both the M/EEG and behavioural measures, then this could be factored in by including time-on-task/trial order as an additional variable in the M/EEG-behaviour analysis (Benwell et al., 2018; Bompas et al., 2015; Macdonald et al., 2011), by regressing out the effect of trial order on the M/EEG and behavioural measures (Wostmann et al., 2018) or by assessing the brain-behaviour relationship over only short timescales (Iemi and Busch, 2018).

Additionally, the current results may have implications for attempts to entrain oscillatory activity through methods such as visual flicker (Ahrens et al., 2015; Capilla et al., 2011; de Graaf et al., 2013; Gulbinaite et al., 2017; Keitel et al., 2018a; Keitel et al., 2014; Mathewson et al., 2012; Notbohm and Herrmann, 2016; Spaak et al., 2014) and non-invasive transcranial brain stimulation (NTBS) (Helfrich et al., 2014; Romei et al., 2016; Thut et al., 2017; Veniero et al., 2017). It has recently been suggested that aligning external stimulation frequency with the endogenous peak alpha frequency may represent an optimal approach to effectively enhance/entrain alpha oscillations (Cecere et al., 2015; Gulbinaite et al., 2017; Romei et al., 2016; Thut et al., 2017; Vossen et al., 2015). Though it remains unclear how essential this alignment of the external and internal oscillator is (Notbohm and Herrmann, 2016; Reato et al., 2013), systematic shifts in peak frequency of the magnitude observed here may be important to consider when attempting to interact with ongoing activity. By contrast, the heterogeneous nature of the sources of alpha activity (Barzegaran et al., 2017; Bollimunta et al., 2008; Buffalo et al., 2011; Capilla et al., 2012; Haegens et al., 2015; Hughes and Crunelli, 2005; Katayama et al., 2011; Keitel and Gross, 2016b; Sadaghiani et al., 2010; Scheeringa et al., 2016) invalidates the assumption that the peak-frequency necessarily represents a single endogenous ‘oscillator’ and hence raises questions regarding the rationale of targeting the peak-frequency for entrainment. Therefore, precise targeting of oscillatory networks will benefit from more refined considerations of the origin and functional roles of the frequency and power content of the EEG signal.

## Acknowledgements

We would like to thank Dr. Mike X. Cohen for EEG analysis scripts and useful discussions regarding interpretation. This work was supported by grants from the Wellcome Trust (grant numbers 098434 and 098433 to GT and JG respectively) and the UK Economic and Social Research Council (grant number ES/I02395X/1 to CSYB).

